# Comparative Genome Analysis Reveals Human Pathogenic Potential of ESBL-*Escherichia coli* Isolated from Swine Microbiomes

**DOI:** 10.1101/2021.07.15.452468

**Authors:** Luria Leslie Founou, Raspail Carrel Founou, Mushal Allam, Arshad Ismail, Sabiha Yusuf Essack

## Abstract

**Background:** Extended-spectrum β-lactamase-producing *E. coli* (ESBL-Ec) harbouring virulence genes in the microbiome of food animals could likely threaten human health but is poorly understood in Cameroon and South Africa. Here, we assessed the resistome, virulome and mobilome of ESBL-Ec isolated from swine microbiomes in these countries, using whole genome sequencing (WGS).

**Materials/methods:** Eleven clonally-related phenotypic ESBL-Ec isolates were subjected to WGS. The isolates were *de novo* assembled using the CLC Genomics Workbench and SPAdes while RAST and PROKKA were used for annotation of the assembled contigs. Prediction of antibiotic resistance genes, virulence factors and plasmids was performed using ResFinder, VirulenceFinder and PlasmidFinder, respectively.

**Results:** Diverse STs were detected with sequence types ST2144 and ST88 predominating and *bla_CTX-M-15_* (55%) as principal ESBL genes. Although the isolates belonged mainly to commensal phylogroups A/B1 (45/28.3%) and C (18.18%), all harboured at least three extraintestinal pathogenic *E. coli* (ExPEC) VFs with one isolate harbouring up to 18 ExPEC VFs.

**Conclusion:** The resistance and pathogenic potential of ESBL-Ec colonizing the gut microbiota of swine in both countries demonstrate the urgent need to implement effective strategies to contain the dissemination of virulent ESBL-Ec through the food chain in Cameroon and South Africa.

## Introduction

Antibiotic resistance (ABR) is a global public health issue that has severe multi-dimensional repercussions not only in humans, where decades of improvements in healthcare outcomes are threatened, but also in the food production industry. The causal relationship between the extensive usage of antibiotics in food animal production and increasing antibiotic resistance in bacteria affecting humans and animals is widely acknowledged. Antibiotics are used for a variety of purposes, including therapeutic and non-therapeutic uses as metaphylactics, prophylactics, and growth-promoters (FAO, 2015, 2016; O’Neill, 2016). Hence, the emergence and spread of ABR across the farm-to-plate continuum puts occupationally-exposed workers (viz. farmers, agricultural practitioners, abattoir workers, food handlers, etc.), their close contacts and consumers at the end of the food chain at risk of contamination or infection by antibiotic resistant bacteria (ARB) and/or antibiotic resistance genes (ARGs). ABR prevention and containment measures should focus not only on humans, but also on animals and their associated environments (Zhang et al., 2017).

*Escherichia coli* is a recognized commensal bacterium of the gastrointestinal tract of humans and animals. The genomic plasticity of *E. coli* strains allows their adaptation to different environments, hence their wide implication in intestinal and extraintestinal infections in both humans and animals worldwide (Sarowska et al., 2019). *E. coli* displays a clonal population structure delineating four main phylogenetic groups (A, B1, B2, and D) with pathogenic strains belonging to group B2 and D while commensal strains belong to group A and B1.

*E. coli* has been suggested as putative reservoir for extended-spectrum β-lactamase (ESBL) resistance and it has been demonstrated that substantial resistance emerges in commensal bacteria especially those present in the gastrointestinal tract where horizontal gene transfer prevails and does occur within and between species and genera (Founou et al, 2016; 2019). ESBL production in *E. coli* is associated with different resistance genes but its most frequently caused by the production of ESBLs encoded by the *bla*_TEM_, *bla*_SHV_ and *bla*_CTX-M_ families, with the latter being the predominant type (Perovic et al., 2014). ESBL-producing *E. coli* have been detected across the animal, human, and environmental interface worldwide, with the emergence of specific clones able to acquire ARGs and virulence genes via mobile genetic elements (MGEs) such as plasmids, transposons, gene cassettes, and other integrative genetic elements (Founou et al., 2016).

Despite the evidence of increasing prevalence of ESBL-producing *E. coli* in food animals, there is still limited information regarding the genetic structure, diversity and relationship of ESBL-*E. coli* isolates from food animals, especially in the pig industry in Sub-Saharan African countries such as Cameroon and South Africa. The objectives of this study were thus to use whole genome sequencing (WGS) and bioinformatics tools to investigate the current population structure, pathogenicity, genetic diversity and resistomes, virulomes and mobilomes, of ESBL-producing *E. coli* isolates from swine microbiomes in Cameroon and South Africa in order to ascertain their pathogenic potential in human health.

## Materials and Methods

### 1. Ethical Considerations

Ethical approvals were obtained from the National Ethics Committee for Research in Human Health of Cameroon (**Ref. 2016/01/684/CE/CNERSH/SP**) as well as from the Biomedical Research Ethics Committee (**Ref. BE365/15**) and Animal Research Ethics Committee (**Ref. AREC/091/015D**) of the University of KwaZulu-Natal. Approvals were additionally obtained from the Cameroonian Ministry of Livestock, Fisheries and Animal Industries (**Ref. 061/L/MINEPIA/SG/DREPIA/CE**) and Ministry of Scientific Research and Innovation (**Ref. 015/MINRESI/B00/C00/C10/C14**). This study was further placed on record with the South African National Department of Agriculture, Forestry and Fisheries [**Ref.** 12/11/1/5 (878)].

### 2. Study Design and Bacterial Isolates

The study sample consisted of eleven putative ESBL-positive *E. coli* isolates that were collected between March and October 2016 as part of a larger study where ESBL-producing *Enterobacterales*, were collected from three abattoirs in Cameroon and two in South Africa (n=2). These isolates originating from nasal (n=6) and rectal swabs (n=5) from healthy pigs processed at abattoirs, were identified as putative, closely related ESBL producers via VITEK 2 system and enterobacterial-repetitive-polymerase chain reaction (ERIC-PCR) analysis, respectively (Founou et al., 2018).

### 3. Identification, ESBL Screening and Antimicrobial Susceptibility Testing

All samples were cultured on MacConkey agar supplemented with 2 mg/L cefotaxime and incubated for 18-24 h at 37°C in normal atmosphere (Founou et al., 2019). All putative ESBL-producers, were phenotypically characterized to the genus level using Gram staining and biochemical tests (catalase and oxidase tests). The isolates were thereafter phenotypically confirmed using the VITEK 2 system.

The VITEK 2 system was further used for ESBL screening along with the double disk synergy test as previously described (Founou et al., 2019). A series of 18 antibiotics encompassed in the Vitek^®^ 2 Gram Negative Susceptibility card (AST-N255) were tested using Vitek^®^ 2 System and (BioMérieux, Marcy l’Etoile, France). Breakpoints of the CLSI guidelines (CLSI, 2016) were used except for the colistin, amoxicillin + clavulanic acid, piperacillin/tazobactam, amikacin for which EUCAST breakpoints (EUCAST, 2016) were considered with *E. coli* ATCC 25922 and *K. pneumoniae* ATCC700603 being used as controls.

### 4. Whole genome sequencing and data analysis

#### 4.1. Purification, Sequencing and Pre-Processing of Genomic Data

GenElute® bacterial genomic DNA kit (Sigma-Aldrich, St. Louis, MO, USA) was used for genomic DNA (gDNA) extraction with the concentration and purity assessed using agarose gel electrophoresis, NanoDrop 8000 spectrophotometer (Thermo Scientific, Waltham, MA, USA), and fluorometric analysis Qubit® (Thermo Scientific, Waltham, MA. USA). Libraries were constructed using the Nextera XT DNA Library Preparation kit (Illumina Inc., San Diego, CA, USA) and subjected to paired-end (2×300 bp) sequencing on an Illumina MiSeq (Illumina Inc., San Diego, CA, USA) machine with 100× coverage. The generated paired-end reads were merged, checked for quality, trimmed, and *de novo* assembled into contigs with the Qiagen CLC Genomics Workbench version 10.1 (CLC, Bio-QIAGEN, Aarhus, Denmark) and SPAdes version 3.11 (Bankevich et al., 2012) to overrule any inherent shortfalls from both assemblers.

#### 4.2. WGS-based molecular typing

WGS data was used to predict *in silico* multi-locus sequence type (MLST) based on the Achtman scheme which considers allelic variation amongst seven housekeeping genes (*adk*, *fum*C, *gyr*B, *icd*, *mdh*, *pur*A and *rec*A) to assign STs (Larsen et al., 2012). In addition to generating an *E. coli* MLST assignment for each isolate, core-genome MLST (cgMLST) was assigned based on a scheme from EnteroBase server (http://enterobase.warwick.ac.uk/species/ecoli) that uses 2 513 loci (Alikhan et al., 2018). EnteroBase was further used for *in silico* phylotype predictions following the Clermont scheme (Clermont et al., 2013) as well as for *fimH* allelic designations (Alikhan et al., 2018). Ribosomal MLST, hierarchical cgMLST clustering, wgMLST were further performed using core genome data in EnteroBase.

#### 4.3. *In Silico* Resistome and Virulome Profiling

ARGs of the *E. coli* genomes were annotated and identified with ResFinder (Zankari et al., 2012) through the bacterial analysis online platform of GoSeqIt tool. The Comprehensive Antibiotic Resistance Database (CARD) platform was concomitantly used for prediction of ARGs and detection of chromosomal mutation (SNPs) in quinolones ARGs *gyr*A, *gyr*B, *par*C and *par*E. The selected threshold and the minimum percentage of the gene length detected were set to 90% identity for a positive match between the reference database and a target genome. VirulenceFinder (Joensen et al., 2014) available from the GoSeqIt tools server along with the comparative pathogenomics platform VFanalyzer from Virulence Factor Database (VFDB) (Liu et al., 2019) were similarly used to predict and annotate virulence factors (VFs), respectively, with a threshold of 90% identity. ExPEC virulence genes including ferric aerobactin receptor (*iutA*), increased serum survival (*iss*); heat-resistant agglutinin (*hra*), temperature sensitive haemagglutinin (*tsh*); P fimbrial adhesin (*papC*); colicin V (*cvaC*) and capsular polysialic acid virulence factor group 2 (*kpsII*) and invasive factor of brain endothelial cells locus A (*ibeA*) of *E. coli* strains responsible for neonatal meningitis in humans were investigated *in silico*. Moreover, the pathogenicity prediction web-server PathogenFinder (Cosentino et al., 2013) was used to predict bacteria pathogenic potential towards human hosts.

#### 4.4. Detection of mobile genetic elements

The RAST SEED viewer (Overbeek et al., 2014) and Artemis Comparison Tool (ACT) were used to identify the presence of transposases and integrons flanking resistance and virulence genes. MGEFInder of the Center for Genomic and Epidemiology was used for the *in-silico* detection of insertion sequences (IS), conjugative genetic elements and transposons allowing investigation of synteny of mobile genetic elements with VFs and antibiotic resistance genes (Durrant et al., 2020). PHAge Search Tool Enhanced Release (PHASTER) server was used for the identification, annotation and visualization of prophage sequences (Zhou et al., 2011). The profile of bacterial plasmid replicons and plasmid incompatibility groups was assessed through PlasmidFinder 2.1 (https://cge.cbs.dtu.dk/services/PlasmidFinder/) and pMLST 2.0 (https://cge.cbs.dtu.dk/services/pMLST/) (Carattoli et al., 2014). Putative CRISP system and Cas cluster were assessed through CRISPRCasFinder (https://crisprcas.i2bc.paris-saclay.fr/CrisprCasFinder/Index).

#### 4.5. Genome visualization and gene annotation

The *de novo* assembled raw reads were annotated using the Rapid Prokaryotic Genome (PROKKA) version 1.12 beta available from EnteroBase, the NCBI Prokaryotic Genome Annotation Pipeline (PGAP) and RAST 2.0 server (http://rast.nmpdr.org)(Aziz et al., 2008) which identified encoding proteins, rRNA and tRNA, assigned functions to the genes and predicted subsystem represented in the genome. The size, GC content, average coverage, length, N50, L50, RNAs and protein coding sequences were obtained for each isolate. The annotated *in silico* predicted proteins and regions were visualized via the JSBrowser of EnteroBase server and RAST. The genomes of the isolates were visualized using the CG Viewer Server (Grant et al., 2012). In addition, the contigs were mapped against the complete genome of *E. coli Ecol_AZ155* (NZ_CP019005.1) for visualization of the genomic organization.

#### 4.6. Comparative phylogenomic analyses

The whole genome phylogenetic relationship was assessed within the study isolates and with a collection of international *E. coli* genomes (*n* = 118) available at the Enterobase *E. coli* genomes repository as of 12 November 2020. The international isolates were closely related *E. coli* strains of similar STs isolated from various sources (humans, livestock and environment).

The *E. coli* isolate YA00194039 (ERS4920643) was used as reference genome with all assembled contigs being aligned against it to determine SNP locations. The phylogeny of the *E. coli* isolates was characterised using the whole genome MLST (wgMLST), core genome MLST (cgMLST) and accessory genome MLST. Phylogenetic relationships among study isolates and between study and international isolates were assessed based on nucleotide alignments of all the genes in the entire genome (wgMLST) and core genome content (core genes that are present in most genomes with ≥95% of nucleotide identity; cgMLST). Moreover, the accessory gene including ARGs, plasmid replicons and phages content was analysed using the Enterobase server which scans the genome against the core ResFinder and PlasmidFinder databases based on a percentage identity of ≥ 90% and coverage of ≥ 70% in order to generate a customized phylogenetic tree to infer the evolutionary relationship within the study isolates and between the study and international isolates. GrapeTree minimum spanning and phylogenetic trees were further built to describe the relatedness among the study isolates and between the study and international isolates. The generated phylogenomic trees were downloaded, and subsequently visualized and edited using MicroReact (www.microreact.org).

## Results

### 1. Baseline characteristics and phenotypic analyses

All the *E. coli* isolates were ESBL producers and high level of resistance to amoxicillin, cefuroxime, cefuroxime axetil, as well as to third (cefotaxime, ceftazidime) and fourth generation (cefepime) cephalosporins were observed. All isolates were resistant to trimethoprim-sulfamethoxazole and susceptible to cefoxitin, ertapenem, meropenem, imipenem and tigecycline. Altogether, three resistance patterns were identified with the pattern AMP.AMC.TZP.CXM.CXM-A.CTX.CAZ.FEP.TMP/SXT (82%) being the most prevalent one. Two isolates displayed multidrug resistance (MDR; resistance to three or more antibiotic families) with one isolate, PN256E8, being resistant to colistin with a minimum inhibitory concentration (MIC) of 8 mg/L. All but two isolates were susceptible to nitrofurantoin. Relevant population data, specimen source, phenotypic and genotypic characteristics for these isolates are summarized in Supplementary Table 1.

### 2. Genomic features

Table 1 and Table S1 depict all the genomic characteristics including length, GC content, N50, coverage, coding sequences, RNAs, rMLST, phylotype, serotype, CRISPR arrays, etc. of the isolates. The genome size of the isolates ranged from 4.5 Mb to 5.3 Mb with a GC content of 50.5 to 50.9 and coverage of 111 to 188.

**Table 1.** Genotypic characteristics of ESBL-producing *E. coli* isolates (Bioproject PRJNA412434)

| Isolate | Accession Number | Country | Sample type | Abattoir | MLST* | Clonal Complex | FimH sub-typing | Phylogroup | Serotype |
| --- | --- | --- | --- | --- | --- | --- | --- | --- | --- |
| **PN017E2II** | VMKK00000000 | Cameroon | Nasal swab | SH001 | 10 | ST10 Cplx | FimH215 | A | O9:H:9 |
| **PR010E3I** | VKOQ00000000 | Cameroon | Rectal swab | SH001 | 44 | ST10 Cplx | FimH54 | A | O89:H4 |
| **PN027E6IIB** | VKOV00000000 | Cameroon | Nasal swab | SH001 | 69 | ST69 Cplx | FimH27 | D | O-:H18 |
| **PR256E1** | VKOS00000000 | South Africa | Rectal swab | SH005 | 88 | ST23 Cplx | FimH1250 | C | O: Uncertain H9 |
| **PN256E2** | VKOT00000000 | South Africa | Nasal swab | SH005 | 88 | ST23 Cplx | FimH1250 | C | O-:H9 |
| **PN027E1II** | VKOW00000000 | Cameroon | Nasal swab | SH001 | 226 | ST226 Cplx | FimH43 | A | O-:H19 |
| **PN091E1II** | VKOU00000000 | Cameroon | Nasal swab | SH002 | 940 | ST448 Cplx | Unknown | B1 | O-:H33 |
| **PN256E8** | QJRZ00000000 | South Africa | Nasal swab | SH005 | 9440 | ST10 Cplx | FimH23 | A | O-:H52 |
| **PR209E1** | VKOO00000000 | South Africa | Rectal swab | SH004 | 2144 | - | FimH87 | B1 | O-:H49 |
| **PR246B1C** | WHRW00000000 | South Africa | Rectal swab | SH004 | 2144 | - | FimH87 | B1 | O-:H49 |
| **PR085E3** | VKOP00000000 | Cameroon | Rectal swab | SH002 | 4450 | - | FimH566 | A | O-:H18 |

### 3. Antimicrobial resistance phenotypes and genotypes

Whole genome-based resistomes corroborated the observed phenotypic profiles. All isolates evidenced relatively similar combinations of resistance genes encoding target modification, antibiotic inactivation, antibiotic efflux pumps and regulators. All *E. coli* isolates harboured *bla_CTX-M_* with bla_CTX-M-15_ (n=6; 54.54%) being the most frequently identified. The *bla_CTX-M-14_* (PR209E1, PR246B1C) was detected in two isolates as was the *bla_CTX-M-1_* (PN256E1, PN256E2). Three isolates simultaneously harboured the *bla_CTX-M-15_* and *bla_TEM-1B_* whilst one isolate, PN256E8 harboured concomitantly the *bla_CTX-M-15_*, *bla_TEM-1B_*, *bla_TEM-141_* and *bla_TEM-206_* (Table 2). Although only two isolates (PR010E3, PN256E8) displayed phenotypic resistance to aminoglycosides, all except two isolates (PR256E1, PN256E2) harboured aminoglycoside resistance genes including str, *aac, aad* and *aph.* Specifically, *aph* was identified in six isolates: *aph(3’’)-Ib* (6/11, 55%) and *aph(6)-1d* (6/11, 55%) genes (Table 2). Eight (73%) isolates harboured different *aad* genes; including *aadA5 (3/11; 27%)*, *aadA1 (3/11; 27%)* and *aadA2(2/11; 18.18)* (Tables 1). Four (36%) isolates harboured *aac* genes including *aac(3)-IIa* (1/11; 9%), and *aac(6)-Ib* (3/11; 27%).

**Table 2.**
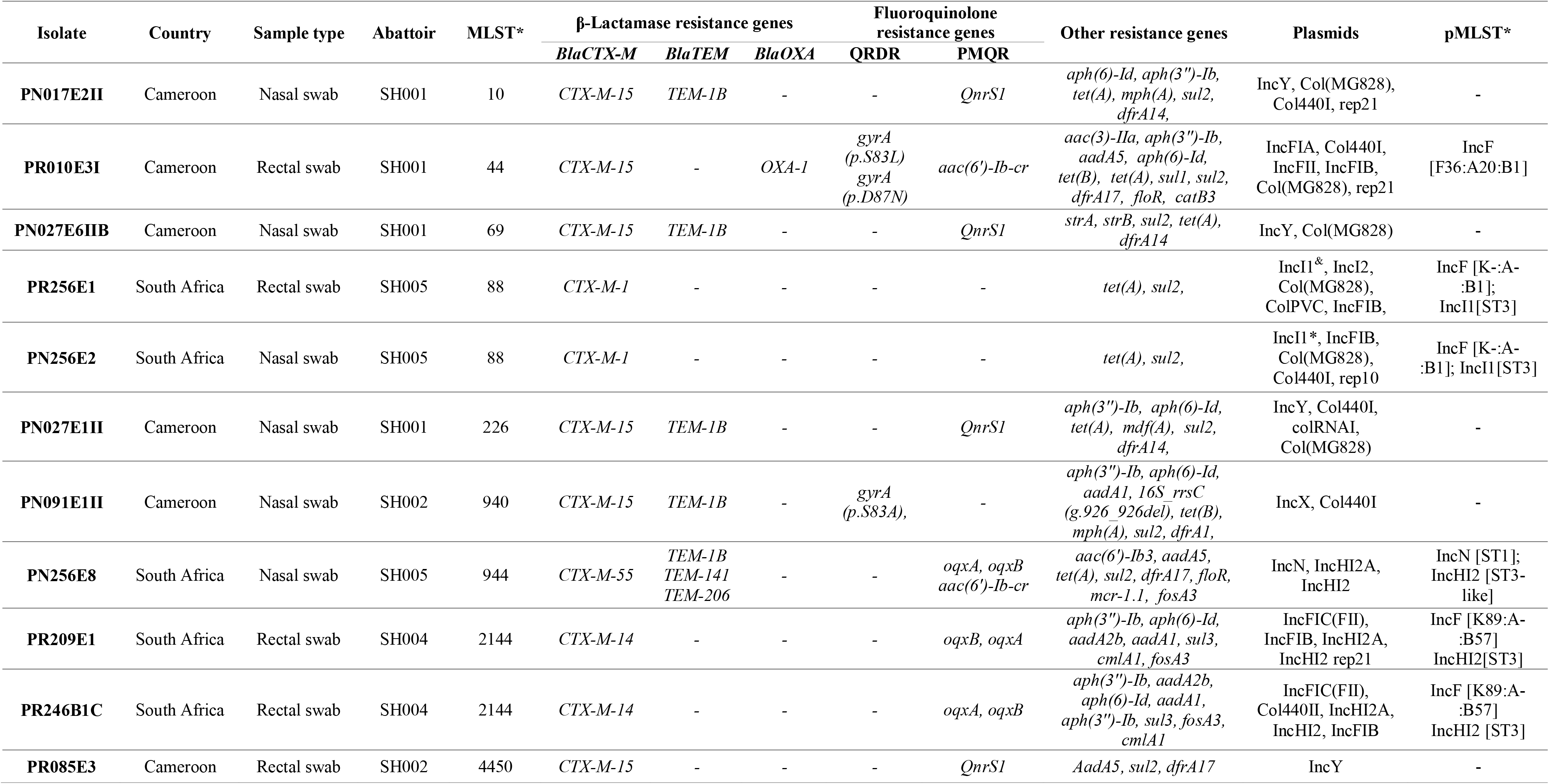
Overview of resistome and mobilome in ESBL-producing *E. coli* isolates.

Several types of plasmid-mediated quinolone resistance (PMQR) genes were identified in the isolates (Table 2). Specifically, the *QnrS1* gene was present in four (36%) isolates, whilst the *aac(6’)Ib-cr* and OqxAB genes were identified in three isolates (27%) each. Mutations in the *gyrA* quinolone resistance-determining region (*QRDR*) genes was observed in two isolates (PR010E3, PN091E1II). *gyrA* had three mutations with two (S**83L**, D**87**N) occurring within PR010E3 and one in PN091E1II (S**83**A) (Table 2). All isolates, except for PN256E8 and PR010E3, for which PMQR gene was identified in both and additionally QRDR in the latter, were susceptible to ciprofloxacin.

All isolates displayed concomitant resistance to trimethoprim and sulfamethoxazole with all harbouring *sul* genes. Specifically, *sul2* gene was identified in nine (82%) isolates alone and in combination with *sul1* gene in one (PR010E3). The rarely evidenced *sul3* gene, was identified in two isolates (PR246B1C, PR209E1). Similarly, the *dfr* gene was identified in 7/11 (64%) isolates, specifically *dfrA17* (n=3), *dfrA14* (n=3) and *dfrA1* (n=1). Diverse permutations of *dfr* and *sul* genes occurred in the isolates. For instance, *sul2* and *dfrA14* were identified in 3/11 (27%) while *sul2* and *dfrA17* were detected in two isolates. One isolate, PR010E3 harboured *sul2*, *sul1* and *dfrA17* concomitantly. In the two isolates harbouring the *sul3* gene, the fosfomycin resistance gene *fosA3* was also detected along with the chloramphenicol resistance gene *cmlA1*. Likewise, in the two isolates harbouring *dfrA17*, the florfenicol resistance gene floR was also detected (Table 2). One isolate (9%) harboured the mcr-1 gene in a full-length copy of a colistin resistance gene showing 100% nucleotide similarity to the reference database sequence (Table 2).

### 4. Whole-genome virulome profiling and pathogenicity

The virulomes of all *E. coli* displayed high level of pathogenicity (Table 3). The ESBL-*E. coli* isolates showed a 93.6% mean probability (P score) of being human pathogens. The pathogenic species with the highest linkage (100% identity) were the *E. coli* APEC O1 (Accession numbers: DQ517526, DQ381420), *E. coli* UMN026 (Accession number: CU928163) and *E. coli* UTI89 (Accession number: CP000243) which are all extraintestinal pathogenic strains in animals (poultry) and humans belonging to the pathogenic phylogroup B2.

**Table 3.**
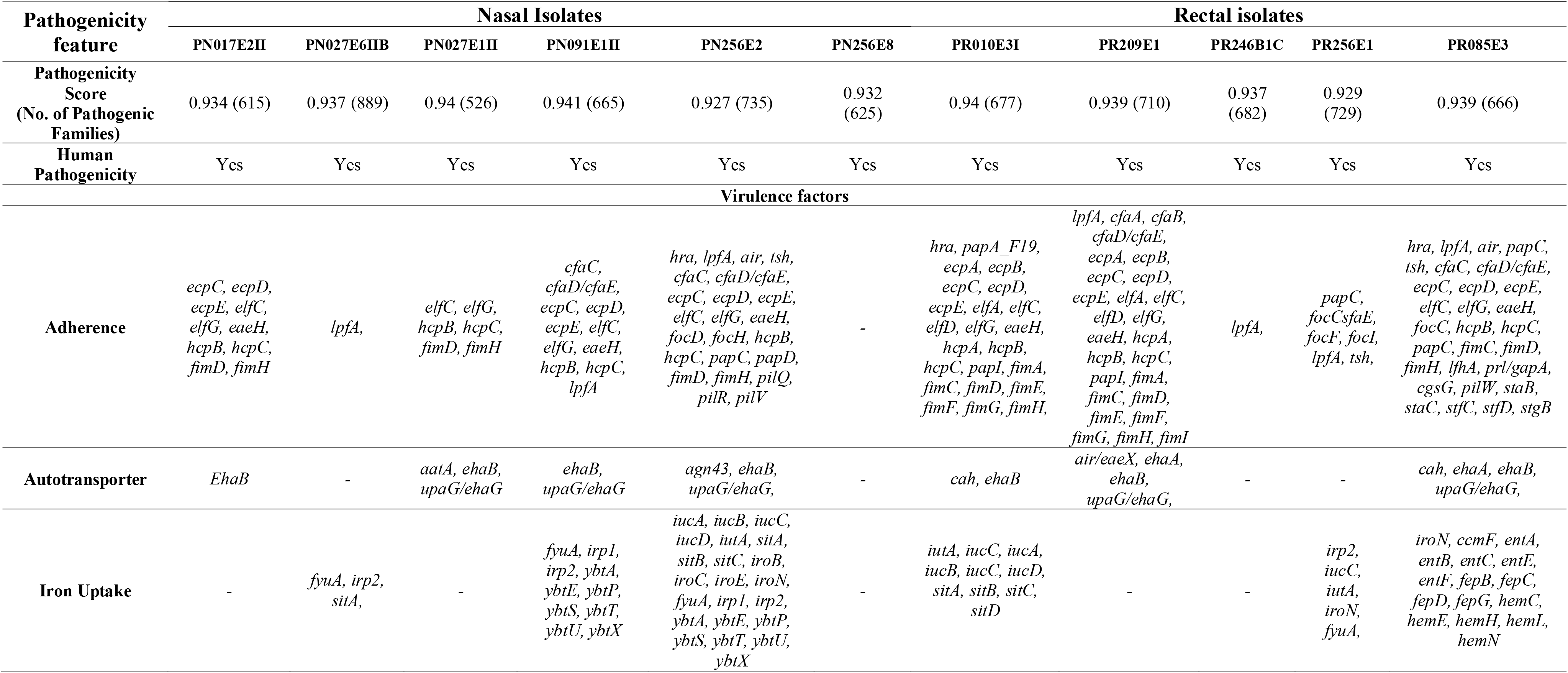
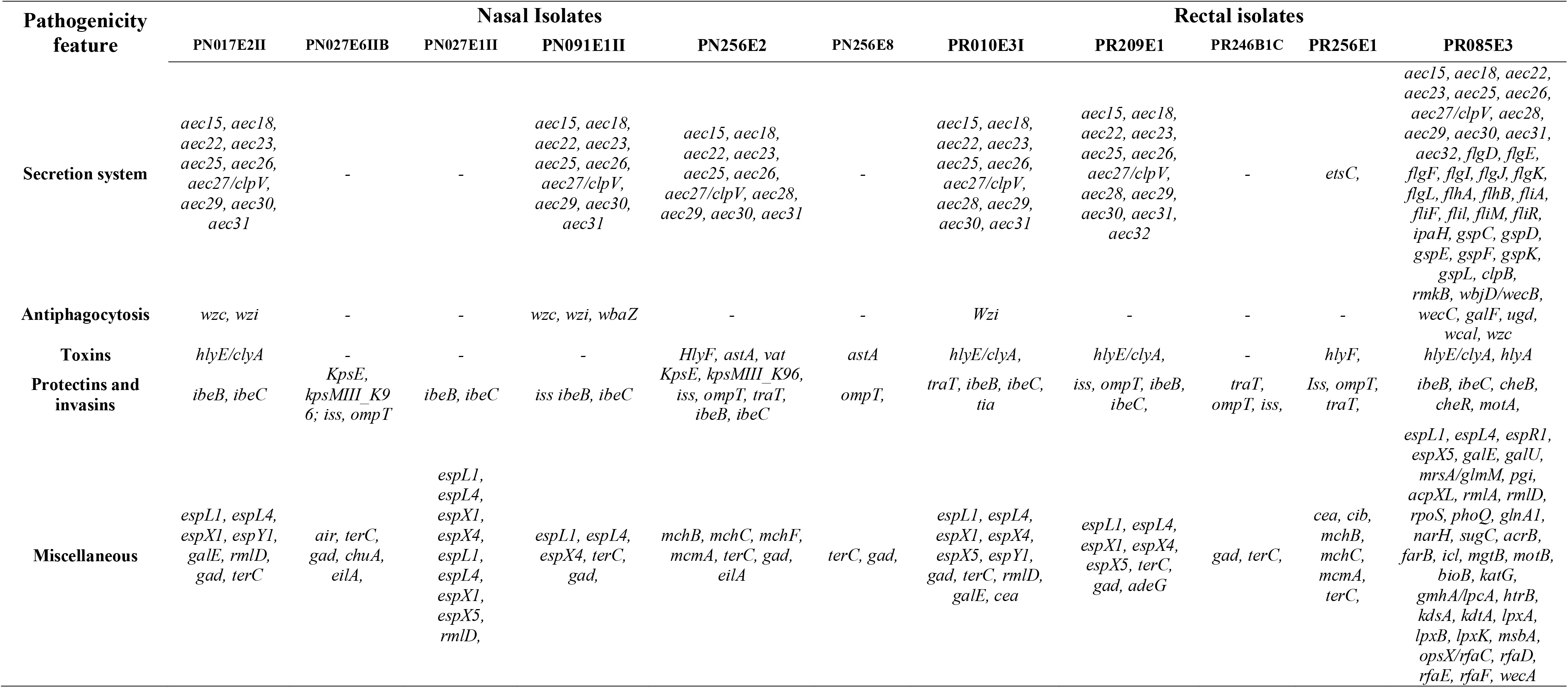
*In silico* identification of human pathogenicity and virulence factors in the ESBL-*E. coli* isolates.

Altogether, over 470 putative virulence factors (VFs) were detected in all isolates. The VFs belonged to major functional categories including: adhesins, toxins, protectins and invasins, iron uptake/siderophores, anti-phagocytosis, secretion systems and autotransporters (Table 3). The isolate PR85E1 harboured the highest number (133) of VFs, followed by the isolates PN256E2 and PR010E3 with 76 and 63 VFs, respectively. All except PR246B1C, PN256E8 and PN027E6II isolates had more than 20 VFs. Analysis of the virulence genotype *fimH* showed that it was present in 64% of the isolates. Among putative VFs, autotransporter adhesin *ehaB*, invasin of brain endothelial cells locus B (*ibeB)* and invasin of brain endothelial cells locus locus C (*ibeC)*, belonging to autotransporter protein and invasins, were the most prevalent (>73%, 8/11) VFs across the isolates (Tables 3-4). The episomal *iss* gene was detected in 55% of isolates whilst *papC,* periplasmic iron-binding protein *sitA (sitA),* siderophore yersiniabactin receptor *(fyuA) genes* were identified in 27% (3/11) isolates. Similarly, the prevalence of protectin genes including the complement resistance protein *traT* was the same as that of iron acquisition genes *iroN and iutA* (Tables 3-4). Interestingly, the avian hemolysin gene F (*hlyF)* that enhance production of outer membrane vesicles (OMVs) and lead to autophagy of eukaryotic cells was detected in one isolate while the hemolysin E (*hlyE*) a pore-forming toxin was observed in 34% (4/11) isolates.

**Table 4.**
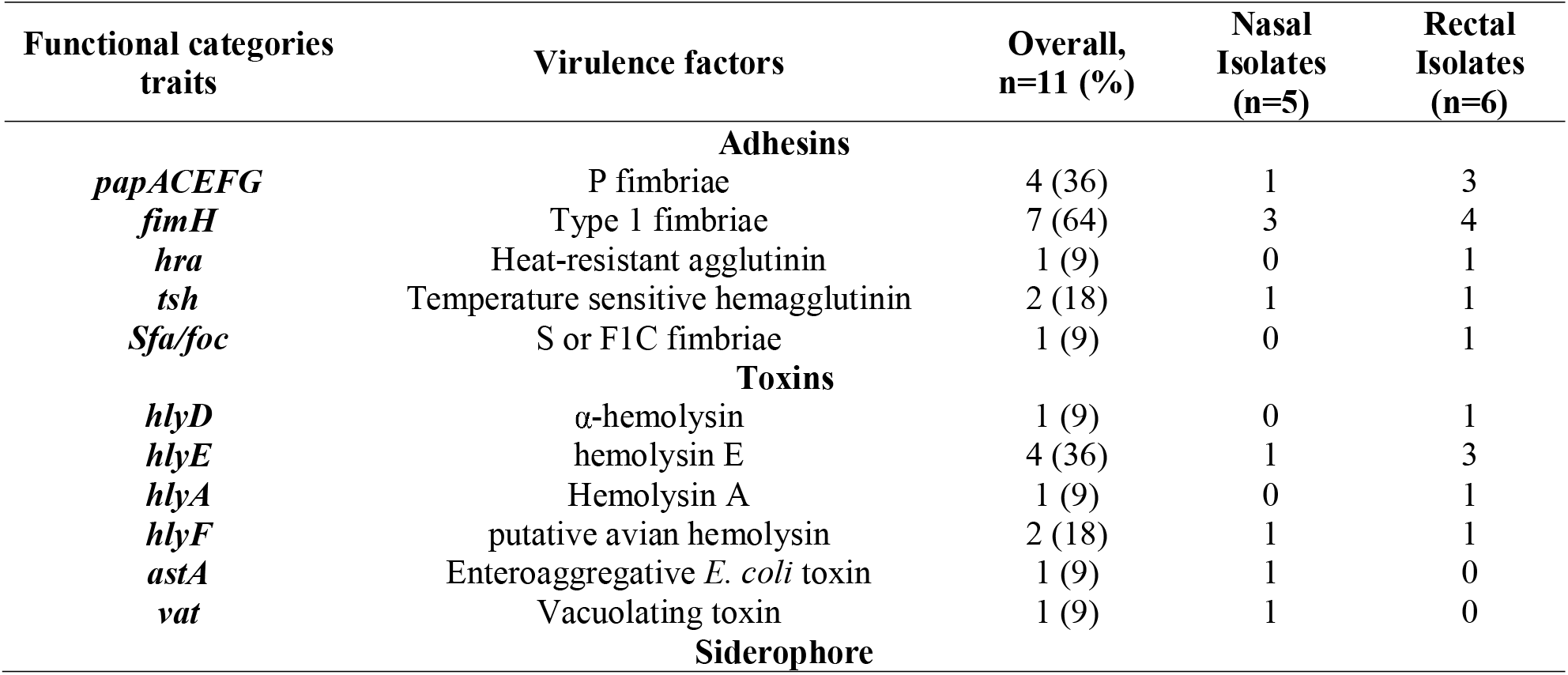
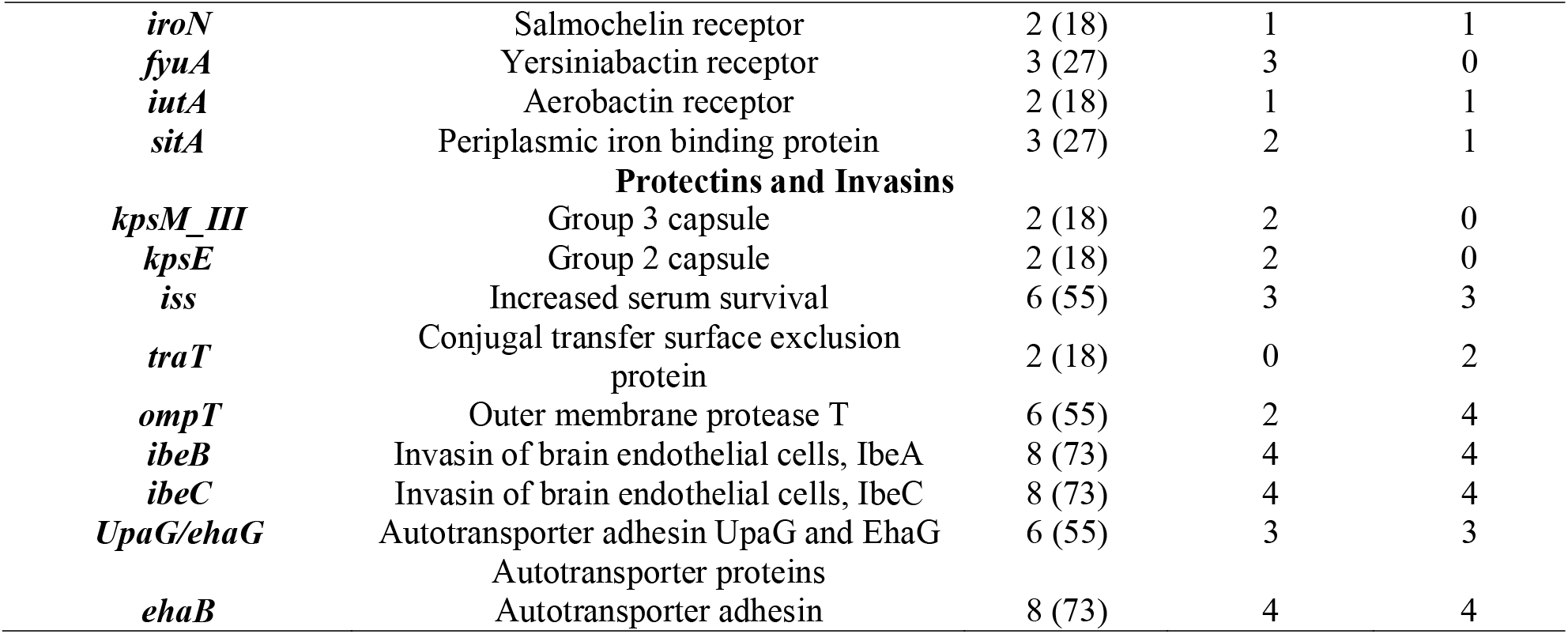
Virulence-associated-traits of nasal and rectal ESBL-*E. coli*.

Our findings showed that rectal *E. coli* isolates harboured more VFs than nasal isolates with prevalence of VFs ranging from 6 to 133 against 4 to 76 in nasal isolates. Putative VFs for invasion, such as the outer membrane protein T (OmpT) and the *traT* genes were most prevalent in rectal than nasal isolates. However, the polysialic acid transport protein group 3 (*KpsMIII)* gene encoding for group 3 capsule, was detected only in nasal isolates as were the unique *vat* and *astA*. Specifically, all *E. coli* isolates harboured at least one ExPEC VF from each of the major functional categories and up to 18 ExPEC VFs (Table 3).

### 5. Phylogenetic groups and multilocus sequence typing, serotyping and phylotyping

Based on *in silico* MLST results, four *E. coli* isolates were assigned to the pandemic ST88 (n=2) and ST2144 (n=2) clones, while the remaining isolates were assigned to six single-locus variants, namely, ST10, ST69, ST226, ST944, ST4450 and ST44. Interestingly, the two *E. coli* ST88 strains, PN256E2 and PR256E1, were detected in nasal and rectal pools of the same pig while the *E. coli* ST2144 were both isolated from two rectal pools of swine originating from the same farm and processed within the same abattoir (SH004).

The majority of the isolates were assigned to commensal phylogroups A (45%), B1 (28%) and C (18%) but one belonged to the virulence phylogroup D (9%). The serotype O-:H49 (18.18%) and O-:H18 (18.18%) were the principal serotypes detected while the *fimH*1250 (18.18%) and *fimH*87 (18.18%) were the predominant *fimH* gene observed.

### 6. Mobile genetic elements

WGS analysis identified 15 different plasmid replicons in all the isolates, which further all harboured multiple plasmid replicons concomitantly. Ten types of incompatibility (Inc) plasmid replicons were identified with different frequencies including IncY, IncFIA, IncFIB (AP001918), IncFIC(FII), IncFII, IncN, IncHI2, IncHI2A, IncI1, IncI2 and IncX (Tables 2 and 5). The majority of isolates (9/11; 82%) harboured the IncF (FII, FIB, FIC, FIA) and IncY (4/11; 36%). Four isolates harbouring the IncF incompatibility group also harboured the IncH (n=2) and IncI (n=2) groups.

**Table 5.**
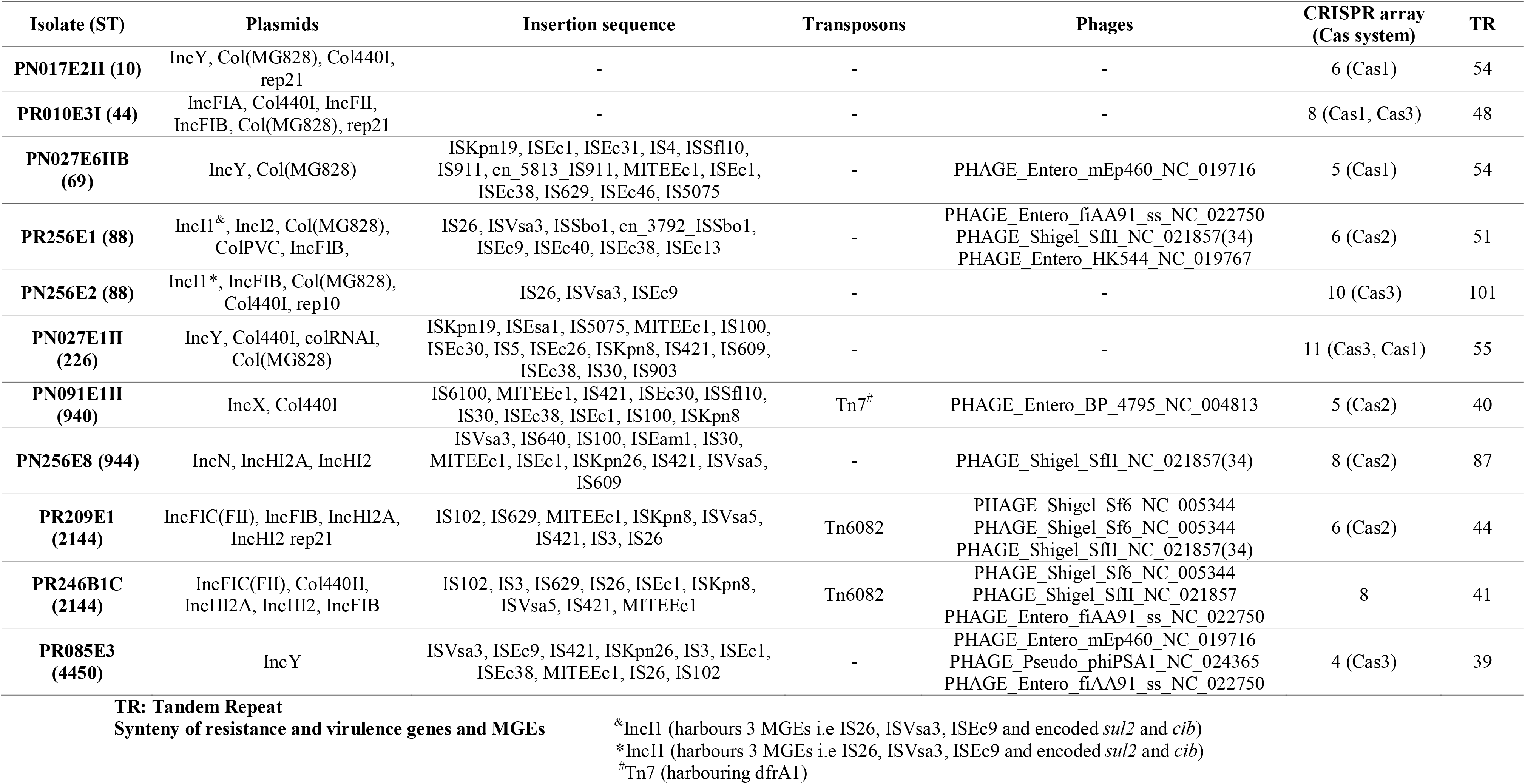
Mobile genetic elements detected in the ESBL-*E. coli* isolates.

**Table 6.**
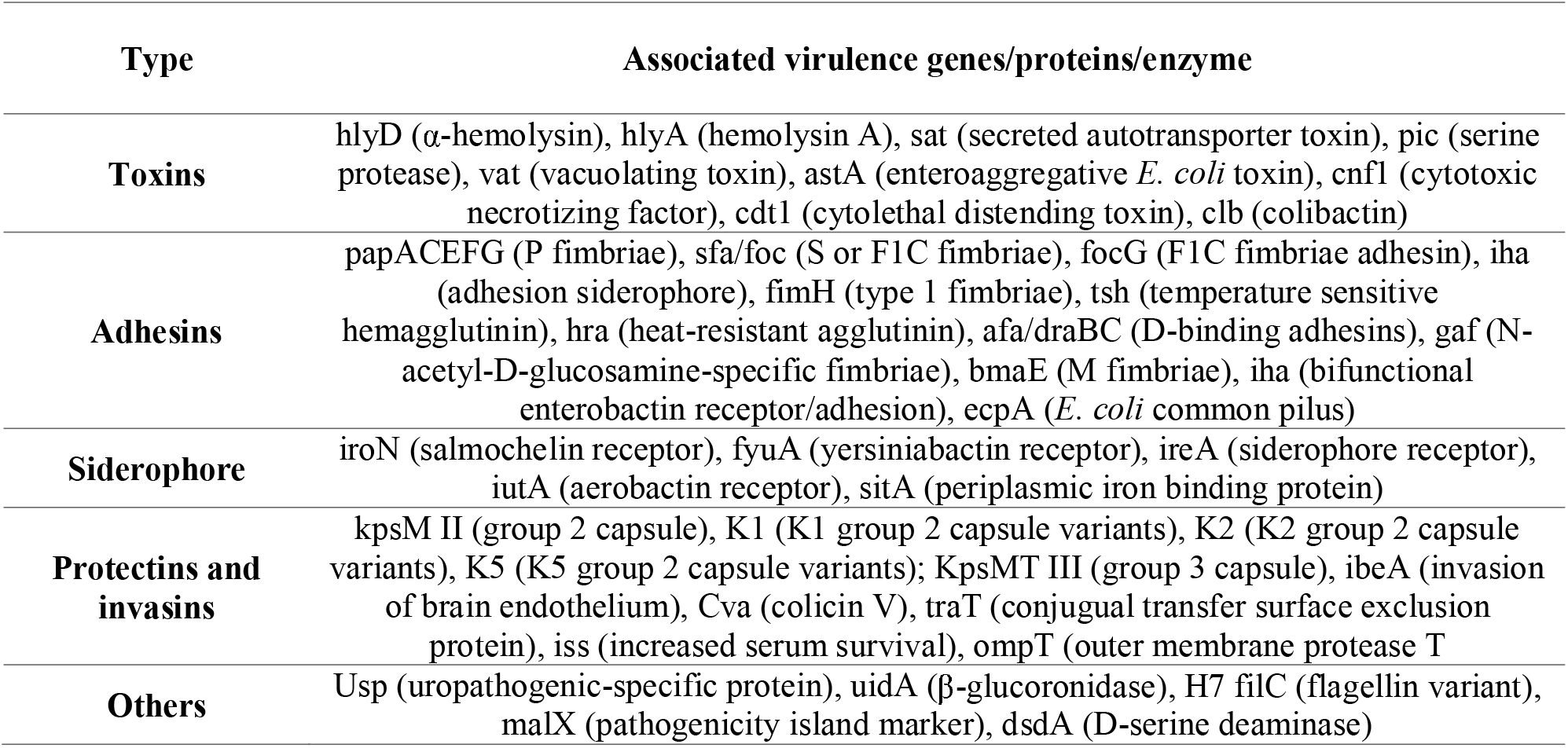
Virulence factors commonly present in ExPEC.

*In silico* plasmid MLST-analyses assigned the IncF plasmid incompatibility group to STs K-:A-:B1 and K89:A-:B57 while IncH and IncI plasmids were assigned to ST3. Additionally, nine (82%) isolates harboured an array of insertion sequences (IS) with IS26, IS421, Isec1 being the most frequent with a 55% prevalence. An array of three IS (IS26, ISVsa3, ISEc9) were harboured on the plasmid IncI that also encoded the sulphonamide resistance gene (*sul2*) and virulence factor (*cib*) in the two *E. coli* ST88 (PR256E1 and PN256E2). Similarly, three isolates harboured transposons (Tn) including Tn6082 (18%) and Tn7 (9%), with the trimethoprim resistance gene *dfrA1* being encoded in transposon Tn7 (Table 5).

### 7. Phylogenetic analysis

The contigs of the ESBL-*E. coli* harbouring the most VFs were mapped against the complete genome of *E. coli* Ecol_AZ155 (NZ_CP019005.1) for visualization of the genomic organisation (Figure 1). Results of comparative genomic analyses revealed specific similarities and dissimilarities, i.e., isolates had similar and dissimilar arrangements of genomic regions towards representative and reference genomes (Figure 1).

**Figure 1.**
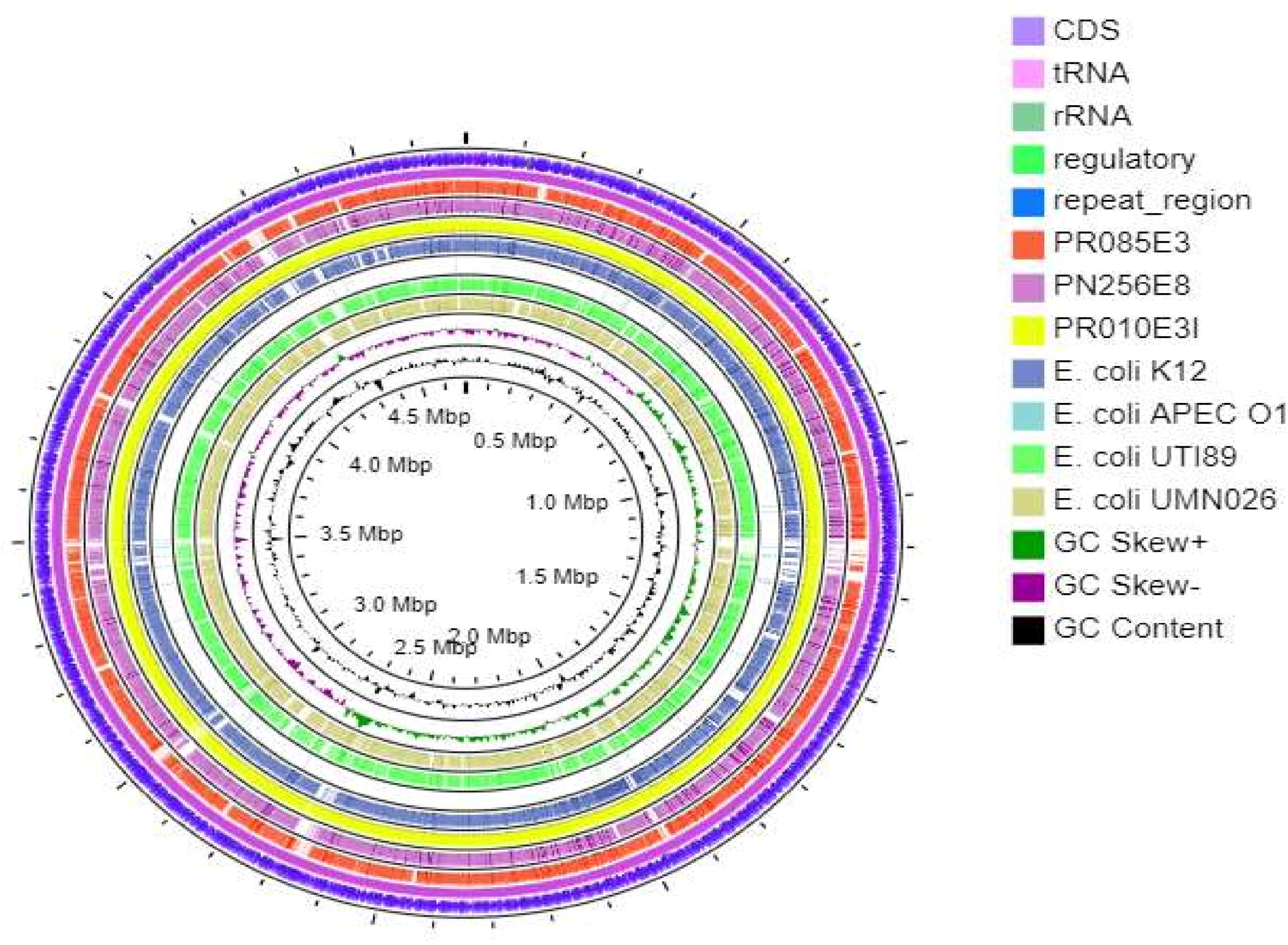
Circular genome representation of selected ESBL-E. *coli* aligned with reference genome and closely related strains. Circular map of selected ESBL-E. *coli* (PR01 0E3, PN256E8 and PR085E3) and comparative alignment with closely related strains (E.coli K12, E.coli APEC_Ol, E.coli UTI89, E. coli UMN026), generated using CGView Server Vl.0. Coloured arrows in the outer ring represent different gene families of the reference genome.A key of the coloured arrows representing different gene families is presented in the inset.The inner coloured circles representing different strains are also listed in the inset. Innermost circles show GC content indicated in black and GC Skew, with green and purple indicating positive and negative values, respectively.

Whole genome phylogenetic analysis grouped the study *E. coli* isolates (n = 11) into two major clusters (Figure 2). The first one grouped six isolates including two *E. coli* ST2144 (PR209E1, PR246B1C), two ST88 strains (PR256E1, PN256E2), one ST940 (PN091E1II) and one ST4450 (PR085E3). The two *E. coli* ST2144 isolates identified from the same abattoir (SH004) in South Africa had 100% identity and shared a close common ancestor with *E. coli* ST940 (PN091E3) and *E. coli* ST4450 (PR085E3) which both originated from one Cameroonian abattoir (SH002). Similarly, the two ST88 isolates (PR256E1 and PN256E2) displayed 100% identity and were found to share common ancestor with the *E. coli* ST2144, ST940 and ST4450. The second clade included four isolates belonging to various STs (i.e., PN256E8: ST9440; PR010E3: ST44; PN017E1: ST10; PN027E1II: ST226) of which *E. coli* ST10 (PN017E1) and ST44 (PR010E3) isolated in the same abattoir Cameroon were closely related and shared common ancestor with *E. coli* ST9440 originating from South Africa.

**Figure 2.**
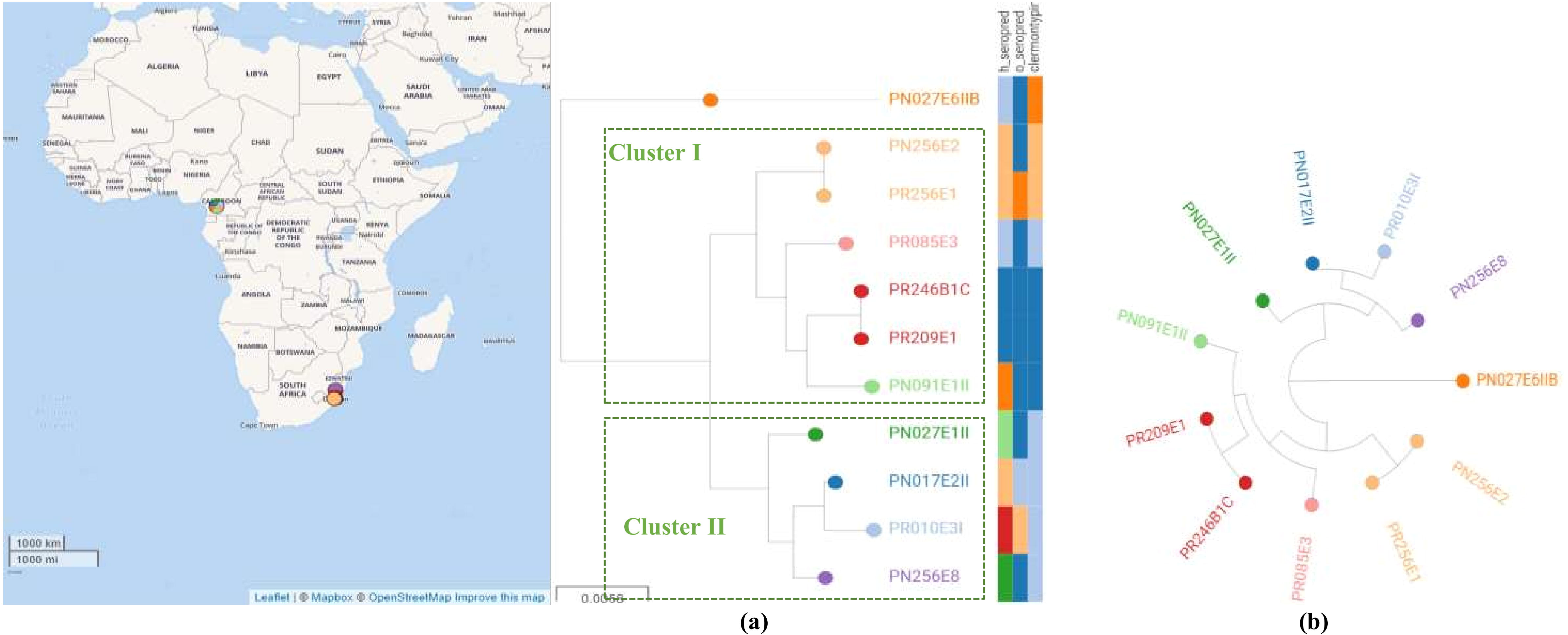
Comparative genome analysis based on the core genome MLST of *Escherichia coli* isolates. Each node represents an isolate, each of which is coloured according to its sequence type, as defined using the core genome. Clusters of isolates belonging to the same sequence cluster are encircled and annotated. Serotype and phylogroup are also indicated via a heatmap (a) Core-genome phylogenetic tree based on comparison of conserved clusters of orthologous genes (COGs). (b) Minimum spanning tree based on cgMLST. Interactive map of geographic locations and genetic attributes can be visualized within Microreact at https://microreact.org/project/tYENaUrCix7jMS7RBrFeBi/09dce9ac

The phylogenomic analyses of the isolates with international strains revealed that they were more closely related to strains from Africa including Kenya, Ghana, Egypt and Morocco than to any other country. None were phylogenetically related to any strain from United States and United Kingdom. The genomes from livestock and humans clustered together at some level. Specifically, PR246B1C and PR209E1 (ST2144), described above to have the same virulome, resistome and mobilome, as well as PN091E1II (ST940), were in the same cluster and shared common ancestors with strains isolated from humans and livestock in Kenya, Uganda, South Africa, Mozambique and Ghana (Figure 3). Similarly, hierarchical clustering analyses provided evidence of genomic relationship between strains originating from livestock and humans with the sole *E. coli* ST10 strain (PN017E2) sharing common ancestor with *E. coli* ST10 isolated from human in Spain and livestock in Luxembourg. Intriguingly, the ST9440 isolate (PN256E8) was closest to a ST10 isolate (Ec416) from livestock in Vietnam. Phylogenetic tree based on accessory genome within the study isolates and between the international isolates revealed similar findings (Figure 4).

**Figure 3.**
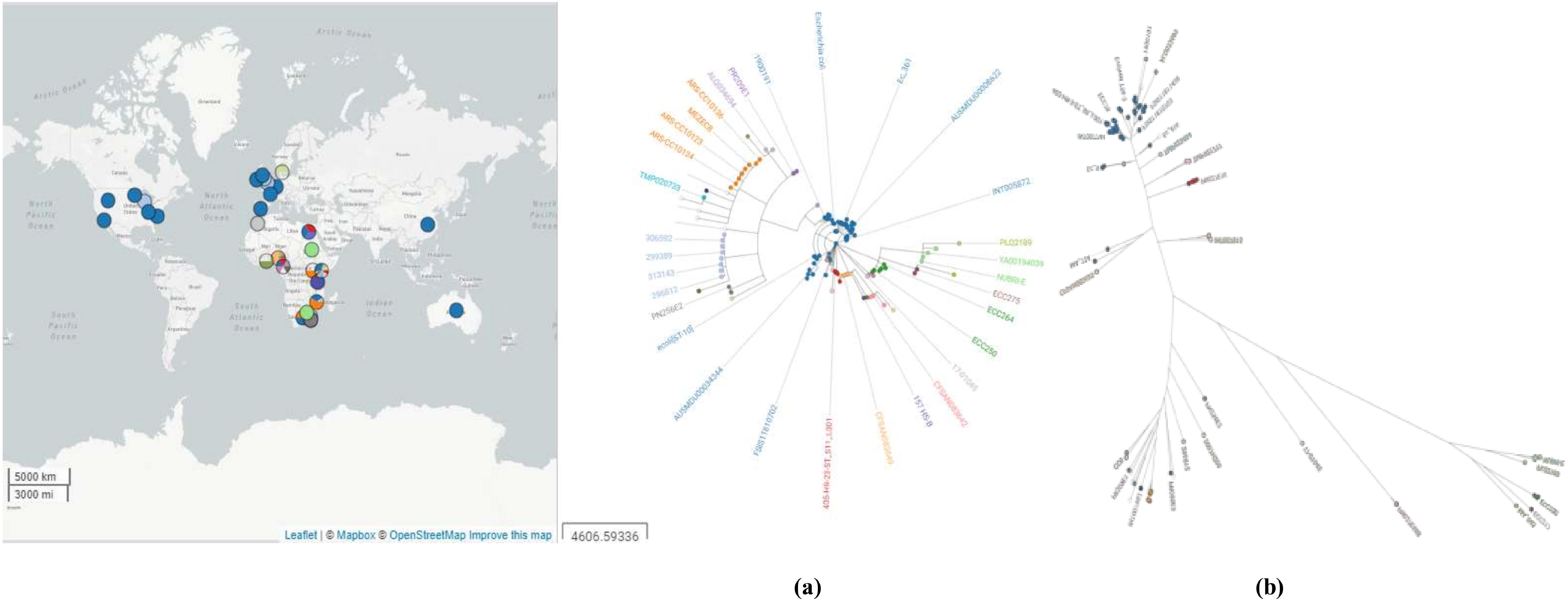
Comparative genome analysis based on the core genome MLST of the ESBL-E. *coli* with international *E. coli* isolates. Isolates of the same STs share the same colour. (a) Minimum spanning tree generated using cgMLST. (b) GrapeTree generated using MSTree V2 tool. Interactive map of geographic locations and genetic attributes can be visualized within Microreact at https://microreact.org/project/2kL6oNgm6VPKnUrrffkUD5/81a2cc99

**Figure 4.**
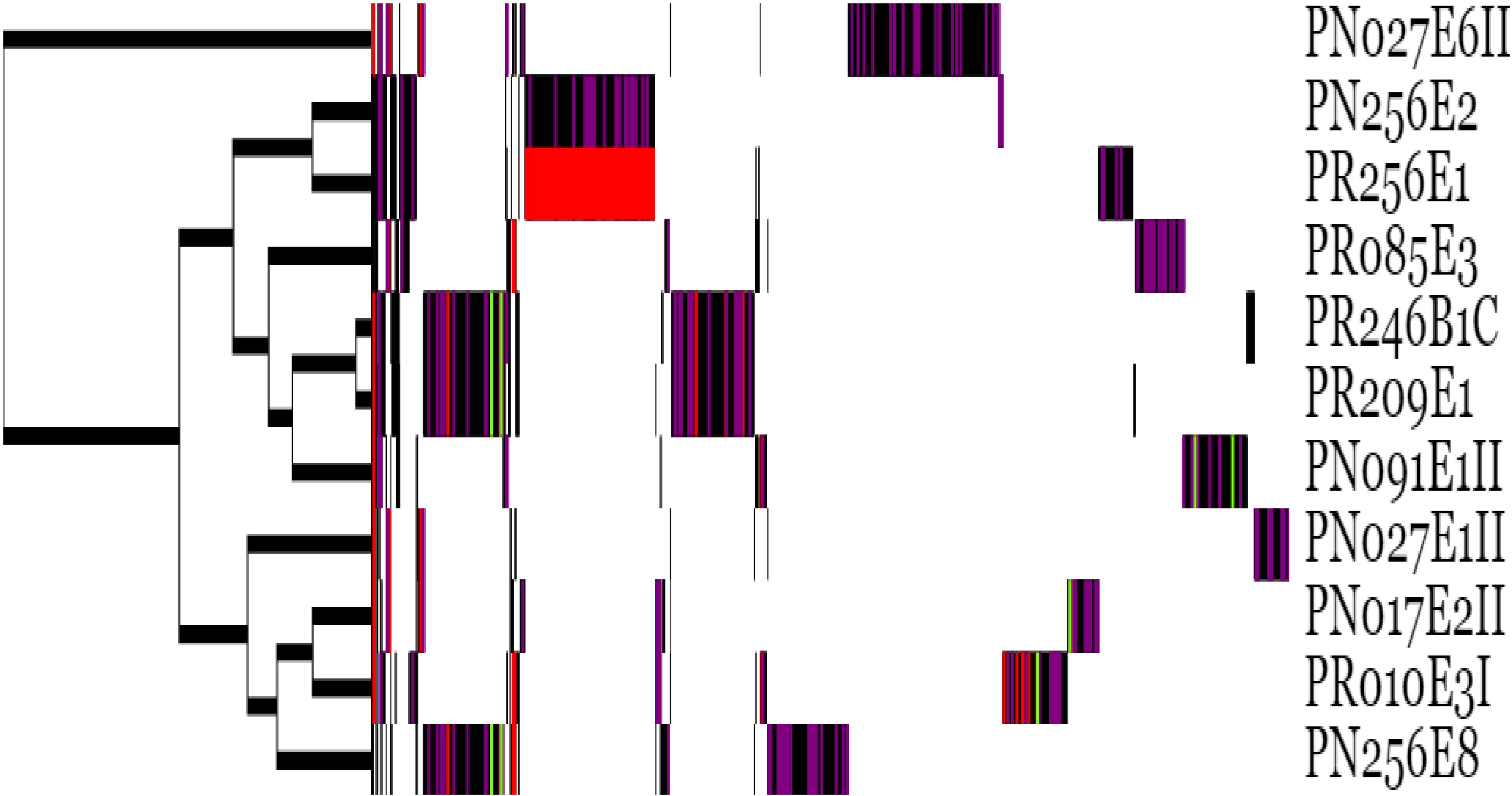
Comparative genome analysis based on the accessory genome of the ESBL-*E. coli* isolates. AMR genes are coloured in red, phages genes (VOGs) in purple and plasmids in are represented in green.

## Discussion

ESBL-*E. coli* is responsible of severe infections worldwide. However, genotypic and pathogenic characteristics of isolates from food-producing animals are not well documented in Cameroon and South Africa. In this study, the population structure of ESBL-*E. coli* isolated from swine microbiomes collected at Cameroonian and South African abattoirs were investigated using WGS. The swine microbiomes from these settings harboured a diverse population of *E. coli* strains that encode an extensive repertoire of resistance genes and plethora of virulence factors, and harboured mainly in IncF, IncH and IncY plasmid replicons.

The antimicrobial resistance data clearly indicates high resistance level of ESBL-*E. coli* isolates to ampicillin, cefuroxime, cefotaxime, ceftazidime and trimethoprim-sulfamethoxazole. Antibiotic use should be limited in agricultural practices and food production systems in order to curtail the emergence of resistance. Each genome of ESBL-*E. coli* isolated from both countries harboured chromosomal and plasmid-mediated ARGs. The ESBL phenotype of the isolates has been confirmed by the presence of ESBL genes (*bla_CTX-M_*, *bla_OXA-1_* and *bla_TEM-1B_*) supporting other contemporary studies which showed that blaCTX-M-15 is the most prevalent ESBL variants among *E. coli,* especially in the highly virulent clone ST131 (Mathers et al., 2015; Rafaï et al., 2015). In fact, Rafai et al. (2015) detected 63.7% of ESBL producers in surgical site infections in Central African Republic. The authors showed that blaCTX-M-15 was present in all isolates along with aac(6’)-Ib-cr. Similar finding were also evidenced by Mbelle et al. (2019) who reported a 70% of blaCTX-M-15 among hospitalized patients in South Africa.

A major observation was the phenotypic multi-drug resistance (MDR) of the isolates PR010E3 and PN256E8 although all isolates, except PR256E1 and PN256E2, harboured concomitantly resistance genes encoding for resistance to aminoglycosides, tetracyclines and fluoroquinolones. This finding is in contrast with data readily available from the literature which suggest that machine or deep learning can be used to predict adequately AMR and MIC of antimicrobial based on genome sequence data (Hendriksen et al., 2019). While transcription analyses could not be undertaken to assess the expression of these genes, these observations led us to posit that the PMQR and QRDR genes as well as aminoglycoside resistance genes present in these isolates might not have been expressed or be silent. Similar discrepancies regarding the phenomes and genomes of isolates harbouring resistance genes but not expressing associated phenotypic resistance were reported elsewhere (Mbelle et al., 2019) and can be further observed in Tables 2 and S2. Our finding further reveals that application of machine or deep learning as well as comparison between phenome and genome are still needed at a large-scale from various environments and sources.

All isolates were seen as human pathogens with over 93% of pathogenicity score and *E. coli* APEC-O1-ColBM (DQ381420), *E. coli* UTI89 (CP000243) and *E. coli* K12 DH10B (CP000948) being the closest related strains. There is increasing evidence that food producing animals and food products or animal origin may contribute to the spread of ExPEC in the community (Hammad et al., 2019). In our study, all ESBL-*E. coli* harboured at least three VFs associated with ExPEC such as *iss*, *iutA*, *traT*, *ompT*, *hlyA*, *iroN*, *papC* and *fimH*. Of great concern is that the isolate PR085EE3 carried over to 100 VFs with the majority having been identified in clinical ExPEC. This suggests that commensal bacteria prevailing in the microbiome of food animals are not only reservoir of resistance genes, but more so, of virulence factors which might be transmitted via HGT and spread to humans through the food chain. It reemphasizes the need to ensure adequate food safety measures throughout the farm-to-plate continuum along with effective infection prevention and control measures in hospitals.

Biofilm formation, essential in colonization and resistance to antibiotics, is an important virulence factor in bacteria. Although biofilm-associated genes have been found to be associated to phylogroup B2 and D strains, biofilm-associated genes such as *fim* and *pap* clusters were detected in our isolates despite being mainly of phylogroup A and B1. We posit that this might again be due to HGT of genetic factors which might have allowed our isolates, though commensals, to acquire these virulence genes horizontally and become putatively virulent in case of extra-intestinal infections. Moreover, the detection of the heat stable enterotoxin 1 (*astA*) gene, encoding for the enteroaggregative *E. coli* heat-stable toxin 1 (EAST1), in the sole ST9440 strain (PN256E8), as well as the avian hemolysin (*hlyF*) in PR256E1 (ST88) gives credence to the fact that commensal *E. coli* prevailing in the gut microbiome have a propensity to acquire various virulence genes which might therefore evolve as progenitor lineages from which heteropathogenic *E. coli* including uropathogenic *E. coli* (UPEC), neonatal meningitis-associated *E. coli* (NMEC) and enteroaggregative *E. coli* (EAEC) strains will emerge.

The majority of commensal *E. coli* strains belong to the phylogenetic groups A and B1, whereas the most common virulent ExPEC are associated with group B2 and D. Our results revealed that the majority of ESBL-*E. coli* isolates belonged to group A (45%) and B1 (28%). Several reports confirmed that phylogroups A and B1 are the leading phylogroups among *E. coli* isolates especially in the gut microbiome (Li et al., 2010; Stoppe et al., 2017). A study from Nigeria showed that 62% of *E. coli* isolates tended towards the commensal phylogroup B1 and A (Olowe et al., 2019). The relationship between phylogenetic groups and ABR has been established previously (Olowe et al., 2019) and studies have shown that the group B2 strains are mainly MDR. However, our study revealed that isolates from other phylogroups such as group A and B1 could also display MDR and high level of virulence certainly as a result of HGT.

Despite the limited sample size (11 *E. coli* isolated from 8 pooled samples), our findings reflect the rich diversity existing within *E. coli* population structure with nine sequence types being detected. The ESBL- *E. coli* isolates were mainly circulating in two clonal lineages since four out of seven isolated strains belong to the ST2144 (n=2) and ST88 (n=2). In addition, the MDR-high-risk clone ST69 and the ST10 were also detected. The ST10 complex, including ST10, commonly associated with spread of CTX-M-1, CTX-M-2 and CTX-M-9 groups, is highly distributed among humans and various livestock species and has been linked with intestinal and extraintestinal infections in several African countries (Rafaï et al., 2015). *E. coli* ST10 was the main ST along with ST131 identified in surgical site infection in Central African Republic (Rafaï et al., 2015). Like other high-risk clones, *E. coli* ST69 possesses biological factors such as *usp*, *ompT*, secreted autotransporter toxin (*sat*) and *iutA* genes corresponding specifically to ST131 (Mathers et al., 2015), that increases bacterial fitness allowing these strains to out-compete other bacterial strains and become the principal part of the bacterial population in the gut (Mathers et al., 2015). The detection of mcr-1 gene in one isolate suggests that colistin resistant *Enterobacterales* are also emerging among food-producing animals in Africa, and demonstrates the urgent need of antimicrobial usage stewardship in food production systems and implementation of effective monitoring programmes to curb the spread of MDR-*E. coli.*

Comparative hierarchical clustering suggested that the majority of our strains belong to two clusters. It further confirmed that ST10 complex is common in African livestock as all our ST10 complex belong to a unique cgMLST cluster containing closely related isolates from Cameroonian (PR010E3, PN027E1II) and South African swine (PN256E8). The comparative phylogenomic analysis further confirms that our ESBL-*E. coli* ST10 demonstrated overlap with ST10 strains isolated from livestock and human populations in Africa (Mozambique, Egypt, Morocco), Asia (Vietnam), Europe (United Kingdom, Denmark, France) and Oceanie (Australia) and displaying the same phylogroup A. Similarly, our ESBL-*E. coli* ST69 share high level of similarity with an UPEC ST69 phylogroup D that was involved in pyelonephritis in France (unpublished data). This gives credence to the hypothesis that commensal *E. coli* of the gut microbiome might be the reservoir of virulence and resistance genes that allow the emergence of hetero-pathogenic *E. coli* strains.

Mobile genetic elements (MGEs) play an essential role in the mobility of ARGs and VFs between different bacterial species. The plasmids of the Inc-family are commonly associated with MDR and VFs in *E. coli* (Rafaï et al., 2015). Ten (91%) of our ESBL-*E. coli* harboured Inc-related replicons with IncF (9/11; 81%) especially the replicon *FIB* (AP001918; 46%) being the leading, followed by IncY (4/11, 36%) and IncH (3/11; 21%) plasmids. Similar plasmid replicons associated with blaCTX-M-group were reported in humans and livestock in Africa and across the world (Patil, Chen, Lian, & Wen, 2019).

Our *E. coli* isolates harboured multiple plasmids belonging to major replicon types and encoding multiple ARGs and VFs. Given the presence of ESBL-*E. coli* in clinically healthy animals and humans, it is likely that the presence of these plasmids could contribute to the long-term persistence of resistance traits in animal and environmental microbiome. Though our study was limited by the isolates numbers and geographic area, our results sufficiently reinforce the need to closely monitor pathogenic and commensal bacteria prevailing in the food production systems on the continent (Patil et al., 2019).

## Conclusion

Our study demonstrates that the population structure of ESBLs-*E. coli* in swine microbiome is highly diverse with the *bla_CTX-M-15_* gene being the leading CTX-M variant. Although the phylogenetic diversity observed precludes any suggestion for clonal dissemination, the resistance and high human pathogenic potential demonstrate the urgent need to implement effective strategies to contain the dissemination of antibiotic resistant bacteria in Cameroon and South Africa. Our study underlines the necessity of long-term and stringent monitoring and molecular studies investigating population structure of commensal and pathogenic bacteria in (food) animals, food products and associated environments, not only to preserve antibiotics for future generations, but also to gain new insights into the diversity, evolutionary history and emergence of ESBL-ExPEC. The Africa Pathogen Genomics Initiative (Africa PGI, https://africacdc.org/download/africa-pathogen-genomics-initiative-factsheet/) recently launched, and aiming at integrating pathogenomics and bioinformatics into outbreak investigations and public health surveillance is expected to advance molecular-based studies and improved infection control and prevention on the continent.

## Acknowledgements

The authors would like to thank all included abattoirs, participants and other involved employees from Cameroon and South Africa for participation and all field work assistance.

Preliminary results from this study were presented in a poster at the 30th European Congress of Clinical Microbiology and Infectious Diseases (ECCMID) in Paris, France, 2020 (abstract 2297).

## Author contributions

**L.L.F** co-conceptualized the study, undertook sample collection, microbiological laboratory and data analyses, prepared tables and figures, interpreted results, performed bioinformatics analysis, and drafted the manuscript. **R.C.F** undertook sample collection, microbiological laboratory analyses, contributed to bioinformatics analysis, vetting of the results and writing of the manuscript. **M.A.** contributed to and vetted bioinformatics analyses. **A.I.** performed whole genome sequencing analysis. **S.Y.E** co-conceptualized the study and undertook critical revision of the manuscript. All authors read and approve the final manuscript.

