## Supplementary tables for "Comparative Genome Analysis Reveals Human Pathogenic Potential of ESBL-*Escherichia coli* Isolated from Swine Microbiomes"

**Table S1. Summary of genotypic features of ESBL-*E. coli***

| Isolate | GC | N50 | L50 | No. Contigs | rMLST | wgMLST | Coverage | Length | Genes | RNA | tRNA | CDS  total | CDS Coding |
| --- | --- | --- | --- | --- | --- | --- | --- | --- | --- | --- | --- | --- | --- |
| PN017E2II | 50.9 | 72955 | 20 | 270 | 2011 | 120277 | 126 | 4 614 573 | 4 626 | 111 | 81 | 4 515 | 4 354 |
| PR010E3I | 50.8 | 69611 | 21 | 226 | 15358 | 128817 | 157 | 4 813 420 | 4 977 | 119 | 80 | 4 858 | 4 635 |
| PN027E6IIB | 50.7 | 117004 | 15 | 179 | 2135 | 120284 | 169 | 4 970 490 | 4 946 | 111 | 81 | 4 835 | 4 696 |
| PR256E1 | 50.5 | 87060 | 20 | 250 | 14767 | 128814 | 131 | 5 312 214 | 5 474 | 105 | 84 | 5 369 | 5 146 |
| PN256E2 | 50.6 | 101064 | 16 | 253 | 14767 | 128816 | 188 | 5 240 610 | 5 426 | 114 | 89 | 5 312 | 5 091 |
| PN027E1II | 50.9 | 57128 | 27 | 257 | 1930 | 128813 | 116 | 4 586 694 | 4 693 | 111 | 80 | 4 582 | 4 350 |
| PN091E1II | 50.8 | 49629 | 27 | 250 | 41587 | 115069 | 131 | 4 822 597 | 4 871 | 121 | 82 | 4 750 | 4 535 |
| PN256E8 | 50.6 | 85550 | 18 | 256 | 32411 | 128815 | 128 | 4 949 938 | 5 315 | 108 | 82 | 5 207 | 5 000 |
| PR209E1 | 50.6 | 76165 | 22 | 219 | 139138 | 114166 | 134 | 5 044 693 | 5 090 | 121 | 84 | 4 969 | 4 851 |
| PR246B1C | 50.5 | 113392 | 13 | 189 | 139138 | 11382 | 171 | 5 071 043 | 4 869 | 108 | 75 | 4 761 | 4 645 |
| PR085E3 | 50.8 | 57886 | 28 | 213 | 38604 | 114162 | 111 | 4 771 008 | 4 793 | 110 | 79 | 4 683 | 4 542 |

ribosomal Multilocus Sequence Type, Whole genome Multilocus Sequence Type, Ribonucleic acid, transfer RNA, Coding sequence

**Table S2. Antimicrobial resistance phenotype of the isolates ESBL-*E. coli***

| Isolate name | β-lactam antibiotics | | | | | | | | | | | Non-β-lactam antibiotics | | | | | | |
| --- | --- | --- | --- | --- | --- | --- | --- | --- | --- | --- | --- | --- | --- | --- | --- | --- | --- | --- |
|  | **AMP** | **AMC** | **TZP** | **CXM** | **CTX** | **CAZ** | **FEP** | **ETP** | **MEM** | **IMP** | **GEN** | | **AN** | **CIP** | **TGC** | **NIT** | **CS** | **SXT** |
| PN017E2II | R | S | S | R | R | R | R | S | S | S | S | | S | S | S | S | S | R |
| PN027E1II | R | S | S | R | R | R | I | S | S | S | S | | S | S | S | S | S | R |
| PN027E6IIB | R | S | S | R | R | R | I | S | S | S | S | | S | S | S | S | S | R |
| PN091E1II | R | R | R | R | R | R | R | S | S | S | S | | S | S | S | S | S | R |
| PR010E3I | R | S | S | R | R | R | I | S | S | S | R | | I | R | S | S | S | R |
| PR085E3 | R | S | S | R | R | R | I | S | S | S | S | | S | S | S | S | S | R |
| PN256E2 | R | S | S | R | I | S | I | S | S | S | S | | S | S | S | S | S | R |
| PN256E8 | R | S | S | R | R | R | I | S | S | S | R | | R | S | S | I | R | R |
| PR209E1 | R | S | S | R | I | S | S | S | S | S | S | | S | S | S | I | S | R |
| PR256E1 | R | S | S | R | R | R | S | S | S | S | S | | S | S | S | I | S | R |
| PR246B1C | R | S | S | R | R | S | S | S | S | S | S | | S | S | S | S | S | R |

AMP: Ampicillin, AMC: Amoxicillin-clavulanic acid; TZP: Piperacillin-tazobactam; CXM: Cefuroxime; CTX: Cefotaxime; CAZ: Ceftazidime; ETP: Ertapenem; MEM: Meropenem; IMP: Imipenem; GEN: Gentamicin; AN: Amikacin; CIP: Ciprofloxacin; TGC: Tigecycline; NIT: Nitrofurantoin; CS: Colistin; TMP/SXT: Trimethoprim-Sulfamethoxazole; S: Susceptible; I: Intermediate; R: Resistant;
